# Venturicidin A affects the mitochondrial membrane potential and induces kDNA loss in *Trypanosoma brucei*

**DOI:** 10.1101/2024.01.12.575445

**Authors:** Dennis Hauser, Marcel Kaiser, Pascal Mäser, Anna Albisetti

## Abstract

Neglected tropical diseases caused by trypanosomatid parasites have devastating health and economic consequences, especially in tropical areas. New drugs or new combination therapies to fight these parasites are urgently needed. Venturicidin A, a macrolide extracted from *Streptomyces*, inhibits the ATP synthase complex of fungi and bacteria. However, its effect on trypanosomatids is not fully understood. In this study, we tested venturicidin A on a panel of trypanosomatid parasites using Alamar Blue assays and found it to be highly active against *Trypanosoma brucei* and *Leishmania donovani*, but much less so against *Trypanosoma evansi*. Using fluorescence microscopy we observed a rapid loss of the mitochondrial membrane potential in *T. brucei* bloodstream forms upon venturicidin A treatment. Additionally, we report the loss of the mitochondrial DNA in approximately 40 to 50% of the treated parasites. We conclude that venturicidin A targets the ATP synthase of *T. brucei*, and we suggest that this macrolide could be a candidate for antitrypanosomatid drug repurposing, drug combinations, or medicinal chemistry programs.

## Introduction

Trypanosomatid parasites cause three of the 20 diseases listed as neglected tropical diseases (NTDs) by the World Health Organization: Human African trypanosomiasis caused by *Trypanosoma brucei (Tb)* ssp., Chagas disease caused by *Trypanosoma cruzi*, and leishmaniasis caused by different species of the genus *Leishmania*.

These vector-borne parasites have life cycles that alternate between the mammalian host and the insect vector. Their energy metabolism depends on the life cycle stage and nutrient availability. For example, procyclic forms of *T. brucei* (PCF) live in a glucose-poor environment within the vector and generate their ATP mainly by oxidative phosphorylation inside their single mitochondrion (1). In contrast, within the mammalian host, the bloodstream forms of *T. brucei* (BSF) are exposed to a glucose-rich environment and therefore produce their ATP by glycolysis within specialized, kinetoplastid-specific organelles called glycosomes (2, 3). The single mitochondrion undergoes extensive remodeling to adapt to different carbon sources and, therefore, strongly differs in shape and function between PCF and BSF: an articulated network in PCF versus a thin tubular mitochondrion devoid of cristae in BSF (4). Another peculiarity of the mitochondrion is the F_0_/F_1_-ATP synthase complex in BSF that operates in the opposite direction compared to the PCF (5, 6). In fact, instead of generating ATP, it hydrolyzes ATP (possibly produced via substrate level phosphorylation (7)), to maintain the mitochondrial membrane potential, which is necessary for the import of proteins and metabolites (5).

ATP synthases are bidirectional machines coupling ATP synthesis or hydrolysis with proton translocation through biological membranes (8). These protein complexes are found in every domain of life and are composed of a membrane-embedded F_0_ region and an F_1_ soluble catalytic head (8, 9). Recently, attention has been brought to the distinct nature of the trypanosomal ATP synthase complex. The ATP synthase complex in *T. brucei* is composed of at least 25 subunits, 15 of which are specific to and conserved among kinetoplastids (4, 10–12).

The chemotherapeutics currently used for the treatment of trypanosomatid infections show issues with toxicity, resistance, high costs, or lack of availability (13–16) (eBioMedicine – The Lancet Discovery Science - editorial https://www.thelancet.com/journals/ebiom/article/PIIS2352-3964(23)00005-1/fulltext) . New treatment options are therefore needed. Repurposing of existing molecules has been successful in the past, as evident by the use of amphotericin B and miltefosine (17–19) to treat leishmaniasis, or fexinidazole (20–22) to treat human African trypanosomiasis caused by *T. b. gambiense*.

Apart from amphotericin B, other macrolide compounds have shown activity against trypanosomatid parasites (23–25). Among these are known inhibitors of the ATP synthase of fungi and bacteria (26, 27): for example, venturicidin A [Fig.1]. Given venturicidin A’s activity against *T. b. brucei* (23) we were interested to determine its potential against other trypanosomatid parasites. We further used *T. b. brucei* as a model to characterize the effect that venturicidin A treatment has on trypanosomatid parasites.

**Fig.1.**
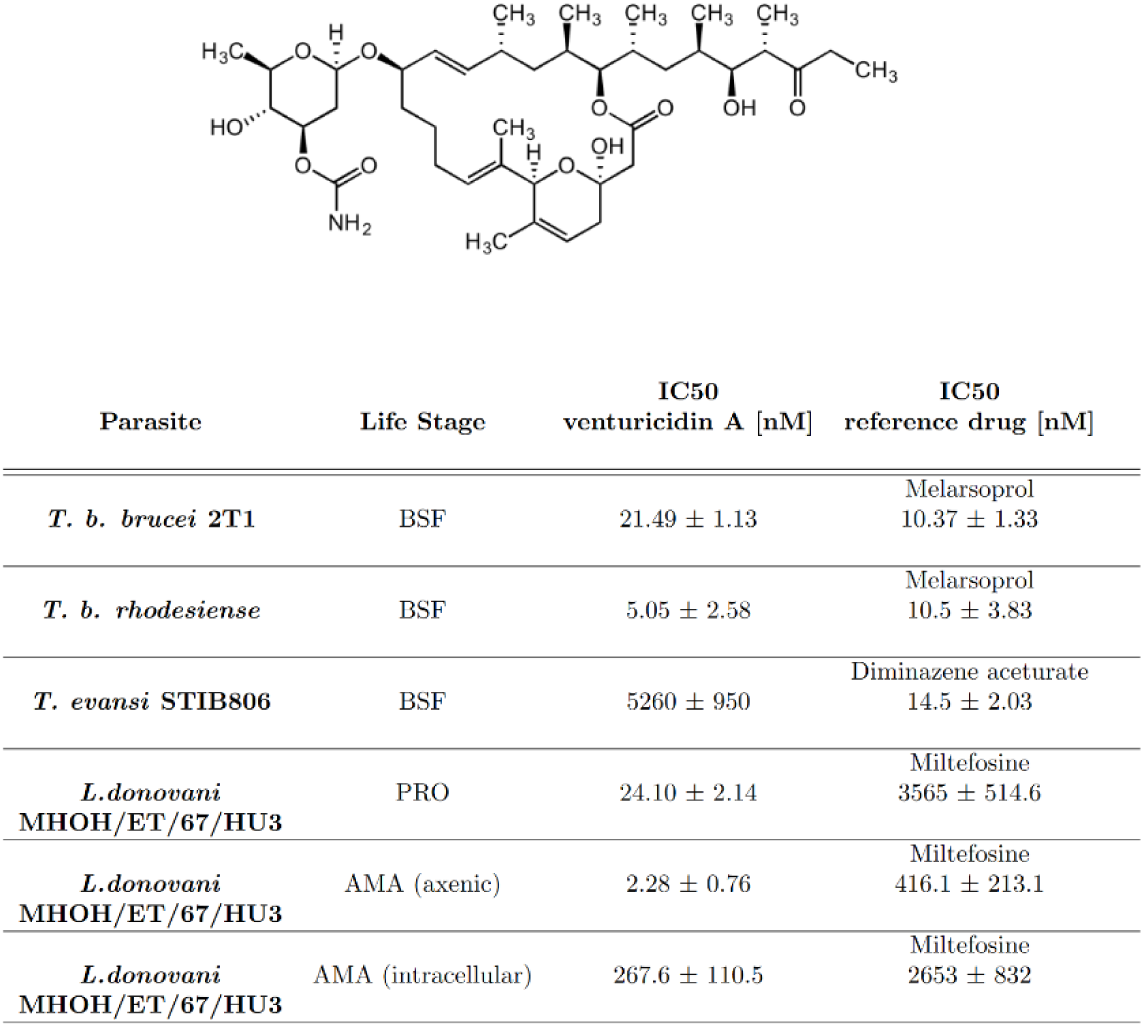
Venturicidin A structure and IC_50_ values. The macrolide venturicidin A (C_41_H_67_NO_11_) (52). 50% inhibitory concentration (IC_50_) of venturicidin A and a reference drug tested against different kinetoplastid parasites. Life stages are abbreviated as follow: BSF: bloodstream form; PRO: promastigote; AMA: amastigote. IC_50_ values are reported in [nM] ± SD; n ≥ 2 (n = 4 for *Tbb*, n = 2 for *Tbr*, *T. evansi*, *L. don* AMA (intracellular), n = 3 for *L. don* PRO, *L. don* AMA (axenic)).

## Results

Venturicidin A [Fig.1] was found to have a selective antitrypanosomal activity *in vitro* (23). We evaluated the *in vitro* efficacy of venturicidin A against bloodstream form *T. b. brucei* 2T1, *T. b. rhodesiense*, *T. evansi*, as well as promastigote and amastigote forms of *Leishmania donovani* using the Alamar Blue assay (28). Venturicidin A showed a 50% inhibitory concentration (IC_50_) against *T. b. brucei* 2T1 of 21.49 nM [Fig.1] and an IC_50_ value of 5 nM against *T. b. rhodesiense* [Fig.1]. Both values are slightly lower than the ones published previously, which were 160 nM for *T. b. brucei* and 720 nM for *T. b. rhodesiense* (23).

Here we report for the first time that venturicidin A acts as a potent inhibitor of the human pathogen *Leishmania donovani*, exerting activity against both the promastigote and amastigote forms. We further show that *Trypanosoma evansi* is less susceptible to venturicidin A than *T. b. brucei*.

Venturicidin A is known to act on the F_0_F_1_-ATP synthase (5, 27, 29, 30). To further investigate its effect on *T. b. brucei*, we treated *T. b. brucei* 2T1 bloodstream forms with 1x IC_50_ (21 nM), 5x IC_50_ (107 nM), or 8x IC_50_ (172 nM) venturicidin A for 24 h and assessed the mitochondrial membrane potential. The treated cells, together with untreated cells as controls, were stained with MitoTracker for 30 min in culture, then fixed and placed onto microscopy slides.

As expected, we observed a clear MitoTracker fluorescence signal in untreated cells, indicative of an intact mitochondrial membrane potential [Fig.2]. In venturicidin A treated cells, independent of the drug concertation used, no MitoTracker signal was visible, indicating the collapse of the mitochondrial membrane potential [Fig.2].

**Fig.2.**
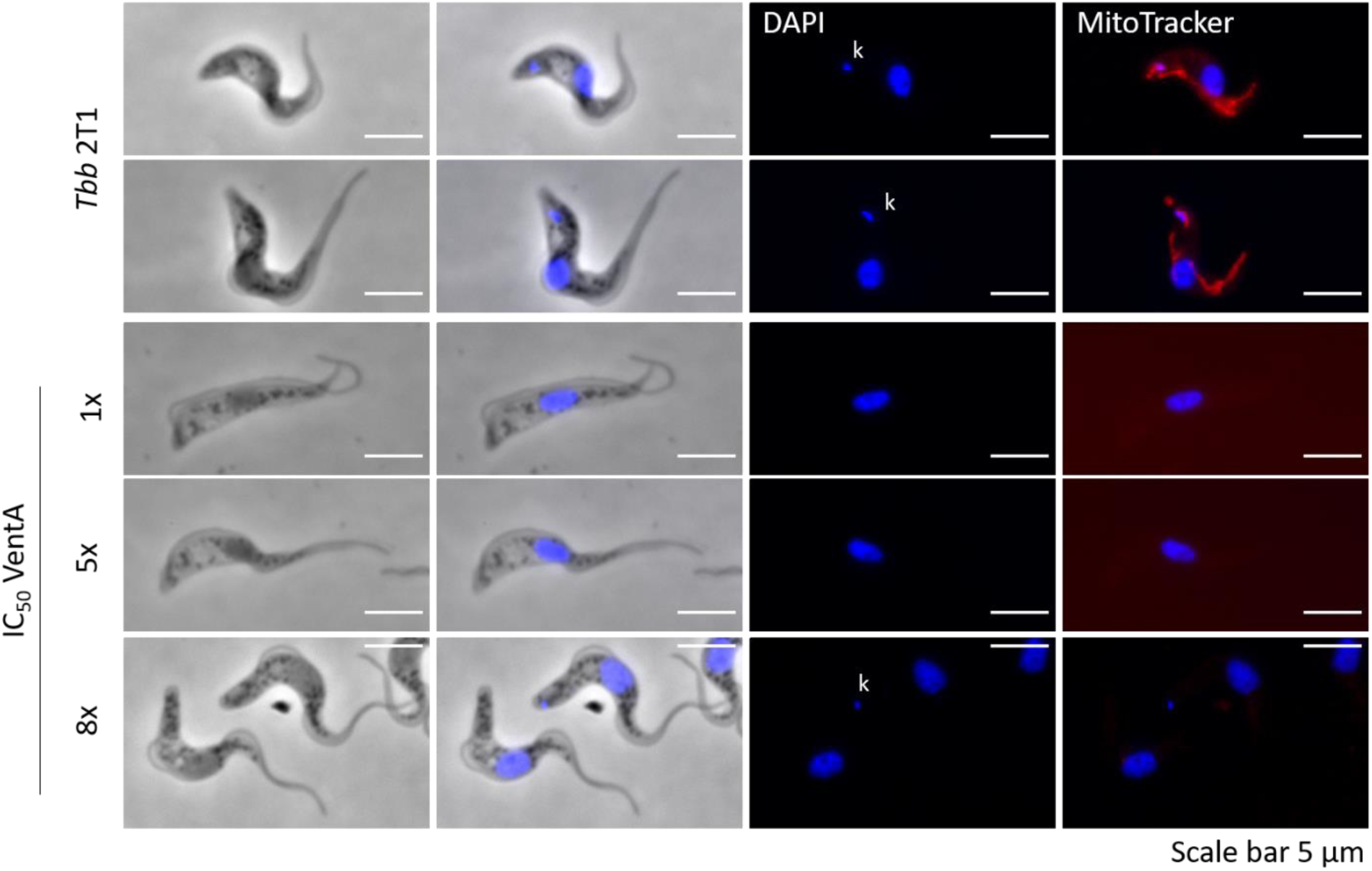
Venturicidin A treatment induces loss of mitochondrial membrane potential. Immunofluorescence of the untreated *Tb brucei* 2T1 cells, and cells treated with 1x, 5x, or 8x IC_50_ venturicidin A (VentA). The cells were stained with DAPI (blue) and MitoTracker dye (red). The kinetoplasts (kDNAs), where present, are marked with (k). Scale bar 5 μm.

Additionally, we noticed that a consistent portion of cells had no detectable mitochondrial DNA, also known as kinetoplast DNA (kDNA) [Fig.2]. To confirm this observation, we treated the parasites with 1x IC_50_ (21 nM), 5x IC_50_ (107 nM), or 8x IC_50_ (172 nM) venturicidin A for 24 h, fixed the cells and stained them with DAPI to mark the nucleus (N) and the kDNA (K), and with an anti-α-tubulin antibody to visualize the cell shape [Fig.3A]. Within a normal population (w/ kDNA = 1K1N, 2K1N, 2K2N), the majority of the cells, around 75-80%, are in the G_1_ phase (1K1N) (31). The percentage of cells that have completed the S-phase and therefore have duplicated both the kDNA and the nucleus (2K2N) and are entering cytokinesis is less than 10 % (31). Compared to the untreated cells, around 40% of the cells treated with either 1x or 5x IC_50_ venturicidin A had lost their kDNA after 24 h, whereas with 8x IC_50_ venturicidin A treatment the percentage of cells without a detectable kDNA reached up to 50% (w/o kDNA = 0K1N) [Fig.3B]. Under all venturicidin A concentrations tested, we also observed dividing cells with an abnormal number of kDNA, such as one kinetoplast but two nuclei (wrong # kDNA = 1K2N) [Fig.3A], in spite of the fact that in the cell cycle of *T. b. brucei*, the kinetoplast divides before the nucleus (32). This possibly indicates how the cells without a detectable kDNA could have originated: a cell with 1K2N divides into a cell with 1K1N and one with 0K1N.

**Fig.3.**
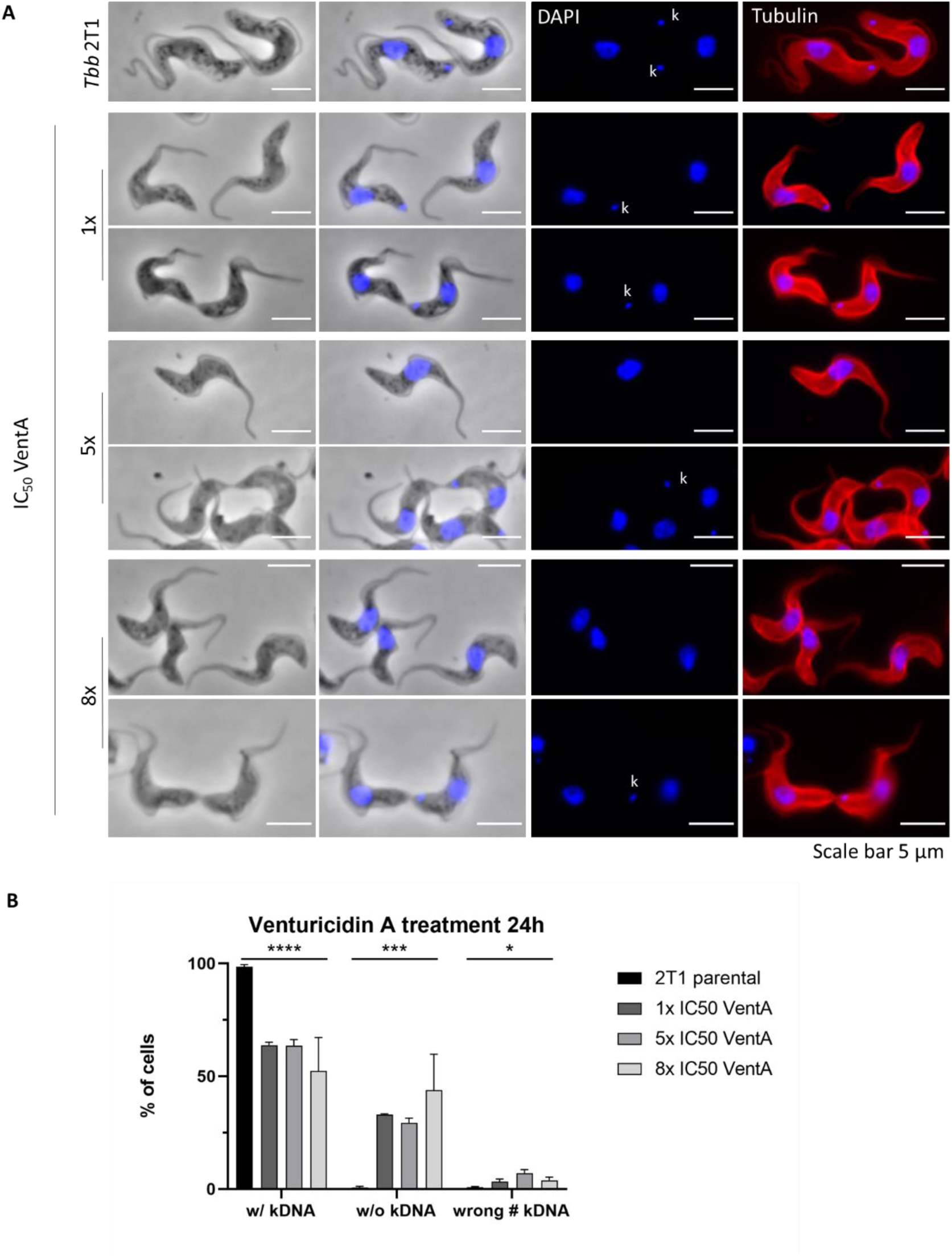
Venturicidin A treatment induces kDNA loss. **(A)** Immunofluorescence of the untreated *Tb brucei* 2T1 cell line, followed by cells treated with 1x, 5x, or 8x IC_50_ of venturicidin A (VentA). The lower panel for each venturicidin A concentration shows dividing cells. The cells were stained with DAPI (blue) as a DNA marker and an anti-α-tubulin antibody (red) to visualize the cell body. The kinetoplasts (kDNAs), where present, are marked with (k). Scale bar 5 μm. **(B)** Percentage of cells showing a normal phenotype (w/ kDNA; e.g. 1K1N, 2K1N, 2K2N), lacking the kDNA (w/o kDNA; 0K1N), or having an abnormal number of kDNAs (wrong # kDNA; i.e.1K2N) in untreated cells, or treated with 1x, 5x, or 8x IC50 venturicidin A for 24 h. Error bars represent standard deviations (n = 3, at least 150 cells per condition). Statistical analysis were done with a t-Test (parametric, unpaired, two-tailed) with 2T1 parental vs pooled 1x – 5x – 8x VentA. P-value **** < 0.0001, *** 0.0002, * 0.0125.

Our results are in agreement with an action of venturicidin A on the ATP synthase resulting in the loss of the mitochondrial membrane potential, and they show for the first time an effect of venturicidin A on the kDNA.

## Discussion

The antitrypanosomal activity of venturicidin A has been known for more than a decade (23). The current experiments resulted in a slightly lower IC_50_ value for venturicidin A against *T. b. brucei* compared to the IC_50_ value reported previously. Both IC_50_ values were determined with the same method (28) but using two different *T. b. brucei* strains: 2T1 in this study and GUTat 3.1 previously. We further observed that venturicidin A’s activity extends to *Leishmania donovani*, the main causative agent of visceral leishmaniasis, as well as the animal pathogen *Trypanosoma evansi*.

Our results in *T. b. brucei* parasites are in agreement with previously published observations regarding the action of venturicidin A on the ATP synthase complex of fungi, bacteria, and trypanosomes (5, 27, 29, 30). In fact, upon venturicidin A treatment, we observed the collapse of the mitochondrial membrane potential, which in *T. b. brucei* bloodstream forms is maintained by the ATP synthase operating in reverse mode. The same effect has been observed previously in *T. b. brucei* with the ATP synthase inhibitor oligomycin (5, 33).

It is the first time (to our knowledge) that loss of kDNA was observed upon venturicidin A treatment. kDNA loss has previously been described as a result of treatment with drugs like the DNA intercalating agents phenanthridine (also known as ethidium bromide) or acriflavine (34–37), but has so far not been reported after treatment with a macrolide.

*Trypanosoma evansi* and *Trypanosoma equiperduum,* responsible for Surra and Dourine, respectively, occur in nature also as dyskinetoplastic forms (36). The introduction of a mutation in the gamma-subunit of the mitochondrial ATP synthase, γ-L262P, compensates for the dyskinetoplasy (absence of kDNA) in *T. b. brucei* (6, 38), without affecting cell viability. In fact, the subsequent loss of gene products encoded in the kDNA is compensated for, by an electrogenic exchange of ADP^3-^ for ATP^4-^ *via* the ATP/ADP carrier, which allows the cells to maintain their mitochondrial membrane potential (38). It is very unlikely that the kDNA loss, observed upon venturicidin A treatment, is the consequence of such a genomic mutation, as the loss of kDNA is already observed within 24 h and because venturicidin A treatment, in contrast to the mutation, induces parasite death. The effect of venturicidin A on naturally occurring dyskinetoplastic trypanosomes requires further attention, especially because induced as well as naturally occurring dyskinetoplastic *T. evansi* have been shown to lose their sensitivity towards the macrolide oligomycin (39). The finding that *T. evansi*, which does not require an active mitochondrion, is less susceptible to venturicidin A than *T. b. brucei* is in agreement with the proposed mode of action, supporting the hypothesis that the trypanocidal activity of venturicidin A is due to collapse of the mitochondrial membrane potential.

The rise of drug resistance is a threat to global health that is increasingly alarming health professionals and scientists. In light of the time- and cost-intensive process of developing new antimicrobial agents, repurposing of already available drugs can be an efficient alternative. Additionally, combination therapies can delay the emergence of drug resistance, as well as lower the dosage of the individual combination partners (40). Both drug repurposing and combination therapies have resulted in new treatment strategies for trypanosomatid infections. A successful example of a combination therapy to treat human African trypanosomiasis is NECT, a combination of nifurtimox and eflornithine (41–43) that is effectively used to treat second stage *T. b. gambiense* infections. One of the most recent examples of successful drug repurposing is fexinidazole, the first orally available treatment against both stages of *T. b. gambiense* infections (22, 44). Venturicidin A is highly active against trypanosomatids and it will be interesting to further study its mode of action. In bacteria, combinations of venturicidin A and aminoglycoside antibiotics have been shown to be synergistic (45). We therefore suggest to also test venturicidin A in combination with known antitrypanosomal drugs.

Understanding the effects of venturicidin A on the trypanosomal ATP synthase complex will provide valuable insight into the biology of trypanosomatid parasites. The introduction of bedaquiline for the treatment of multidrug-resistant *Mycobacterium tuberculosis* infections (46) illustrates that the ATP synthase can be successfully targeted in order to fight microbial infections (CDC Guidelines: https://www.cdc.gov/mmwr/preview/mmwrhtml/rr6209a1.htm). Therefore, we suggest that it is worth studying the ATP synthase complex as a potential drug target in the context of trypanosomatid parasites.

## Methods

### Strains and cultivation

*T. b. brucei* bloodstream form 2T1 were cultured at 37°C with 5% CO_2_ in HMI-9 medium (47), supplemented with 10% foetal calf serum (FCS – BioConcept Ldt 2-01F00-I).

*T. b. rhodesiense* were cultured at 37°C with 5% CO_2_ in HMI-9 medium (47), supplemented with 15% horse serum (self-extracted – Pferdemetzgerei Bürgi - Allschwil).

*T. evansi* STIB806 BSF were cultured at 37°C with 5% CO_2_ in BMEM Minimum Essential Medium (50 µl) supplemented with 25 mM HEPES, 1g/l additional glucose, 1% MEM non-essential amino acids (100x), 0.2 mM 2-mercaptoethanol, 1mM Na-pyruvate and 15% horse serum (48).

*L. donovani* MHOM/ET/67/HU3 promastigotes were cultured at 27°C in SM:SDM79 medium (49, 50), supplemented with 10% FCS. *L. donovani* axenic amastigotes were cultured at 37°C with 5% CO_2_ in SM medium pH 5.4 supplemented with 15% FCS (49).

Peritoneal mouse macrophages (PMM) from CD-1 mice were cultured at 37°C with 5% CO_2_ in RPMI 1640 medium with 10% FCS.

### Drug Assay

Venturicidin A (Santa Cruz sc-202380A) IC_50_ values were measured by Alamar Blue assay as described previously in (28), and for intracellular assays as described in (51); n ≥ 2 (*Tbb* 2T1 BSF n=4, *Tbr* BSF n=2, *T.evansi* BSF, n=2, *Ldon* PRO n=3, *Ldon* AMA axenic n=3, *Ldon* AMA intracellular n=2). The fluorescence was quantified with a SpectraMax reader (Molecular Devices) and SoftMax Pro 5.4.6 Software. Graphs and statistics were made with GraphPad Prism 8. Two-way ANOVA Dunnett’s multiple comparison test between 2T1 parental and 1x – 5x – 8x. P-value **** < 0.0001.

### Immunofluorescence and imaging

For imaging, parasites were incubated with 500 nM MitoTracker (Invitrogen M7513) for 30 min at 37°C with 5% CO_2_, before proceeding with slide preparation. Immunofluorescence analysis were performed as in (31). Anti-α-tubulin DM1α antibody (sc-32293) was used as primary antibody, followed by anti-mouse AF596 (Invitrogen A11012) or anti-mouse AF488 (Invitrogen A11001). Slides were mounted with Vectashield® antifade mounting medium (H-1000). Slides were imaged with a Leica DM5000B fluorescent microscope, equipped with a Leica K5 camera, images were acquired with LAS x software v3.7. Images were processed with ImageJ Fiji. At least 150 cells per condition, in triplicates, were counted.

## Acknowledgment

We would like to thank Sonja Märki-Keller and Romina Rocchetti for *T. b. rhodesiense*, intracellular amastigote *L. donovani,* and *T. evansi* assays, and the Swiss National Science Foundation for financial support (grant No. 310030_185163).

